# An ice-bucket challenge: investigating ice algae physiology in laboratory microcosms

**DOI:** 10.64898/2026.07.10.737583

**Authors:** Margaret L. Baker, Emma Forss, Regina Kolzenburg, Sinéad Collins, Sven A. Kranz

**Affiliations:** Department of Biosciences, Rice University, Houston TX 77005, USA; Institute of Ecology and Evolution, School of Biological Sciences, University of Edinburgh, Edinburgh, EH9 3FL; Umeå Marine Sciences Centre, Umeå University, Norrbyn, Sweden

**Keywords:** sea ice, ice tank, photophysiology, environmental gradient, temperature and salinity covariation

## Abstract

John Raven pioneered the field of algae ecophysiology, advancing our understanding of cellular resource economics, carbon acquisition, and energy allocation. His work laid the foundation for investigating integrative physiology, linking growth-survival trade-offs across diverse environments. The sea ice habitat provides an excellent framework to continue the research John championed. With steep temperature-salinity gradients, algae survival requires a shift in physiology that we are only beginning to understand. We developed two small scale, reproducible icecosms to investigate physiological changes associated with incorporation into sea ice and survival potential post-melt. *Fragilariopsis cylindrus* and *Nitzschia frigida*, known for their association with the ice environment, and *Porosira glacialis*, known for its association with the ice edge, were used to mechanistically link physical properties with algal physiology and post-melt survival. We observe incorporation into the ice of *F. cylindrus* and *N. frigida* alongside vertical photophysiological profiles of *F. cylindrus* revealing inhospitable conditions in the top compared to the bottom layers of ice. *N. frigida* and *P. glacialis* remain viable within the ice and retain the capacity to seed populations following melt. Our results establish icecosms as experimental framework to investigate ecophysiological responses of sea ice algae and provide a foundation toward ecological and evolutionary questions.

## 1. Introduction

Algae represent some of the most ecologically resilient organisms on Earth, occupying almost all environments around our planet. As primary producers representing the base of the aquatic food web, they provide the essential energy source for almost all aquatic ecosystems. These organisms are adapted to specific niches spanning broad environmental gradients, from subzero polar seas to hot springs and from nutrient-rich (eutrophic) to nutrient-poor (oligotrophic) systems. Assessing how these organisms, and especially microalgae, persist and thrive under extreme physical and chemical conditions was a central question in John Ravens’ work (Raven and Geider 1988). Over many decades, John framed algal physiology in terms of cell resource economics, focusing on how cells acquire inorganic carbon and nutrients, allocate energy, and adjust their metabolic budget as light, temperature, and other environmental conditions change (Barton et al. 2020; Beardall and Raven 2004; Raven 1991). His studies and syntheses on photosynthetic performance, CO_2_-concentrating mechanisms, carbon and nutrient acquisition, and on associated energetic costs of the metabolic processes to the cell’s energetic budget provided the basis for describing microalgae as organisms with finely tuned strategies shaped by selection around the world, but especially in challenging habitats.

Polar environments serve as a particularly interesting region to further test his ideas, as they impose multiple interacting environmental extremes, including strong seasonality, low temperatures, and unusual physical and chemical conditions. The sea ice environment in particular offers steep environmental gradients in temperature, salinity, and irradiance. When sea ice forms, the pelagic phytoplankton community is scavenged from the water column into brine channels and pockets throughout the ice. Brine channels within the ice matrix are highly variable, subject to extreme temperature (-20 °C to -1.8 °C; (Petrich and Eicken 2017)), salinity (>200 PSU; (Arrigo et al. 2010)), and irradiance (0 to 1000 µmol photons m^-2^s^-1^; (Galindo et al. 2014)). Upper sea ice layers can approach ambient air temperatures, while insulation by the ice itself and the warmer underlying water column can produce steep vertical temperature gradients (Arrigo 2014). Similarly, irradiance in the upper sea ice layers can approach direct solar levels, but strong scattering within the ice leads to rapid attenuation with depth, reflected in extinction coefficients several-fold higher than in clear water (Light et al. 2008; Perovich 1996). Throughout the ice column, these brine channels represent micro-habitats with steep gradients in which ice algae are partitioned into (Arrigo and Thomas 2004).

Studying microalgal growth and physiology in sea ice is currently limited by the logistical difficulties of field-based sampling (Thomas and Dieckmann 2002), the prohibitive costs and low throughput/replication of existing laboratory ice-growth systems (Kennedy et al. 2012; Thomas et al. 2021; Yoshida et al. 2020, 2021), and the potential inaccuracies of extrapolating data from purely pelagic aquatic environments (e.g. bottle experiments with high salinities) to the unique sea-ice matrix. To address these limitations, we have developed a replicable, low-cost sea-ice microcosm system designed for the study of algal ecology and physiology and used this platform to better understand ecological and physiological processes of three selected ice-algae species within the ice and/or during the critical post-melt transition. This system allows for the testing of the ‘resource economics’ framework. Particularly, it allows us to quantify how microalgae adjust their metabolic budgets, balancing photosynthesis with the metabolic costs which come with cellular acclimations to extreme environments such as sub-zero temperatures, hypersalinity, and light fluctuations characteristic of the polar brine channel environment.

Freezing temperature and high salinity place constraints on cellular function and are likely to have fitness repercussions. Changes in growth rates are usually a combination of direct effects and physiological adjustments, such as the production of antifreeze proteins (Kirst 1990; Thomas and Dieckmann 2002; Uhlig 2011) or elevated expression of core proteins such as Rubisco (Young et al. 2015). While it was generally assumed that temperature has no effect on the light reaction of photosynthesis (Falkowski and Raven 2007; Raven and Geider 1988), freezing temperatures reduce membrane fluidity (Morgan-Kiss et al. 2006), affecting enzyme turnover (Morgan-Kiss et al. 2006; Raven 1990; Rehder et al. 2025; Young et al. 2015) within the light reaction pathways and beyond. In his work, John Raven often emphasized that the Q10 temperature coefficient, the factor by which a reaction rate increases with a 10 °C rise, is much higher for enzymatic ‘dark’ reactions than for physical ‘light’ or transport processes (Raven and Geider 1988). Consequently, sub-zero temperatures inhibit enzymatic kinetics to such an extent that nutrient acquisition significantly outpaces the cell’s capacity for biosynthetic processing (Priscu and Sullivan 1998). Thus, the effective photosynthetic performance and the effectiveness of electron flow through the photosynthetic electron transport chain (PETC) is strongly reduced (Mock and Junge 2007). Other physiological acclimations to freezing temperature include the upregulation of non-photochemical quenching strategies to dissipate excess energy from the photosystem (Ralph et al. 2005). Algae may also reduce the area available for light harvesting (σPSII) to reduce energy absorbed (Dubinsky et al. 1986) to avoid light excitation in the first place. As freezing temperature appears to increase processes protecting against light stress rather than facilitating efficient light use, low temperatures may ultimately affect the cell’s ability to utilize light, even at reduced irradiance (Maxwell et al. 1994). In addition to specific adaptations of the photosystem to temperature stress, sea ice algae must maintain osmotic plasticity to manage variable salinity within the ice (Ewert and Deming 2013; Ralph et al. 2005). As salinity increases, sea ice algae are known to demonstrate reduced photosynthetic efficiency, capacity, and growth rate. Changes in photosynthetic efficiency and rates of electron transport under high salinity can limit the reduction of the primary electron acceptors and the PQ pool (Kirst 1990; Satoh et al. 1983). Additionally, sea ice algae may alter their osmolyte concentration to acclimate to such dramatic changes in salinity (Ralph et al. 2007).

The sea-ice and brine channel environment raises the kind of questions of integrative physiology John Raven championed, such as which acclimatory responses are most effective, what are their energetic and elemental costs, and how do they trade off against growth and survival across the seasonal trajectory of ice formation, consolidation, and melt. Inspired by John Raven’s guiding research principles and his conceptual clarity within the field of algal ecophysiology, we aim to continue his effort by examining how sea-ice algal physiology is regulated when microorganisms are embedded within the ice matrix. Specifically, we focus on how constraints imposed by the brine environment influence the coupling between light use efficiency, photoprotective capacity, and carbon fixation. By viewing sea-ice algae through Raven’s resource-allocation framework, we treat ice not just as an extreme temperature regime, but as an extreme, complex and highly dynamic environment where unique physiology has evolved. We offer a unique tool for replicated and controlled studies that allow us to connect mechanistic physiology to ecological function of how microalgae maintain productivity, sustain carbon fixation, and support polar food webs while also identifying traits that could control resilience or vulnerability as sea-ice extent, duration, and internal structure continue to change in a warming climate.

To address distinct ecological objectives, we designed two specialized icecosm systems. In this initial study, we utilized these modular systems to determine if key polar ice-associated diatom species exhibit preferential incorporation into the ice-matrix during ice formation, and to assess their subsequent growth dynamics when released from distinct (upper vs. lower) vertical sections during melt-out. One larger icecosm was used to investigate *Fragilariopsis cylindrus* distribution and photophysiology within an ice matrix, while smaller icecosms were used to demonstrate the post-melt growth potential of two additional diatom species, *Nitzschia frigida* and *Porosira glacialis*.

## 2. Methods

### 2.1. Construction of icecosms

Two specialized icecosm systems were designed: a Square-Icecosm (SIC) and a Mini-Icecosm (MIC). The SIC was designed specifically to quantify energy dynamics and physiological stress across varying light and salinity gradients, utilizing a high-volume capacity tank that allows for high-resolution temporal sampling (up to 50 ice cores from a single unit). Conversely, the MIC was developed as a compact modular platform to facilitate the simultaneous observation of multiple species or biological replicates in parallel. In this study, we utilized these modular MIC systems to determine if key ice-associated diatom species exhibit preferential incorporation during ice formation, and to assess their subsequent growth dynamics when released from distinct (upper vs. lower) vertical sections during melt-out. Both icecosms share a very similar construction - an inner container housed in an insulated outer layer, both with an open top. An aluminum heat exchanger with internal flow channel (e.g. Peltier water cooling plate) was installed at the bottom outside of the inner container connected to a water pump and a water reservoir to adjust the temperature within the heat exchanger (Figure 1). Cooling of the system to below freezing was achieved within a walk-in freezer at -20 °C or in a household chest freezer maintained at a temperature of ∼ -10.4 ± 1.2 °C throughout the experiment (Table 1). The combination of exterior insulation, the heat exchange on the bottom of the tub and an internal submerged impeller pump (e.g. tmc Reef Pump Compact 500 (MIC) or SOALR/DC WATER PUMP IP68 (SIC)) ensured ice formation from surface-down. This setup ensures that ice forms at a controlled rate from the top down and creates a consistent temperature gradient. Our icecosms varied in size, with the SIC being 40 L using a polyethylene tub (30 x 46 x 30 cm depth, width, height) and series of five temperature probes (Coralife Digital Aquarium Thermometer) anchored within the SIC at 5 (chamber), 12, 16, 20, and 23 cm (water) from the top of the tank to monitor temperature at each sampling point, ranging from the external freezer chamber to the underlying Aquil* seawater. The MICs consisted of 5L cylindrical containers (VWR Beaker without Handle, 76019-276; radius 10.1 cm, height 26.1 cm). The MIC used a recirculating glycerol:water bath (3:10) initially set to −1 °C but adjusted throughout the experiment to control the ice buildup. The temperature in the MIC was measured by drilling holes ∼3-4 cm horizontally into the ice and inserting a temperature probe directly into the ice. The salinity of the core was measured by removing cores with an ice screw and placing a core, either in its entirety or cut into sections, into a sterile falcon tube to melt.

**Figure 1:**
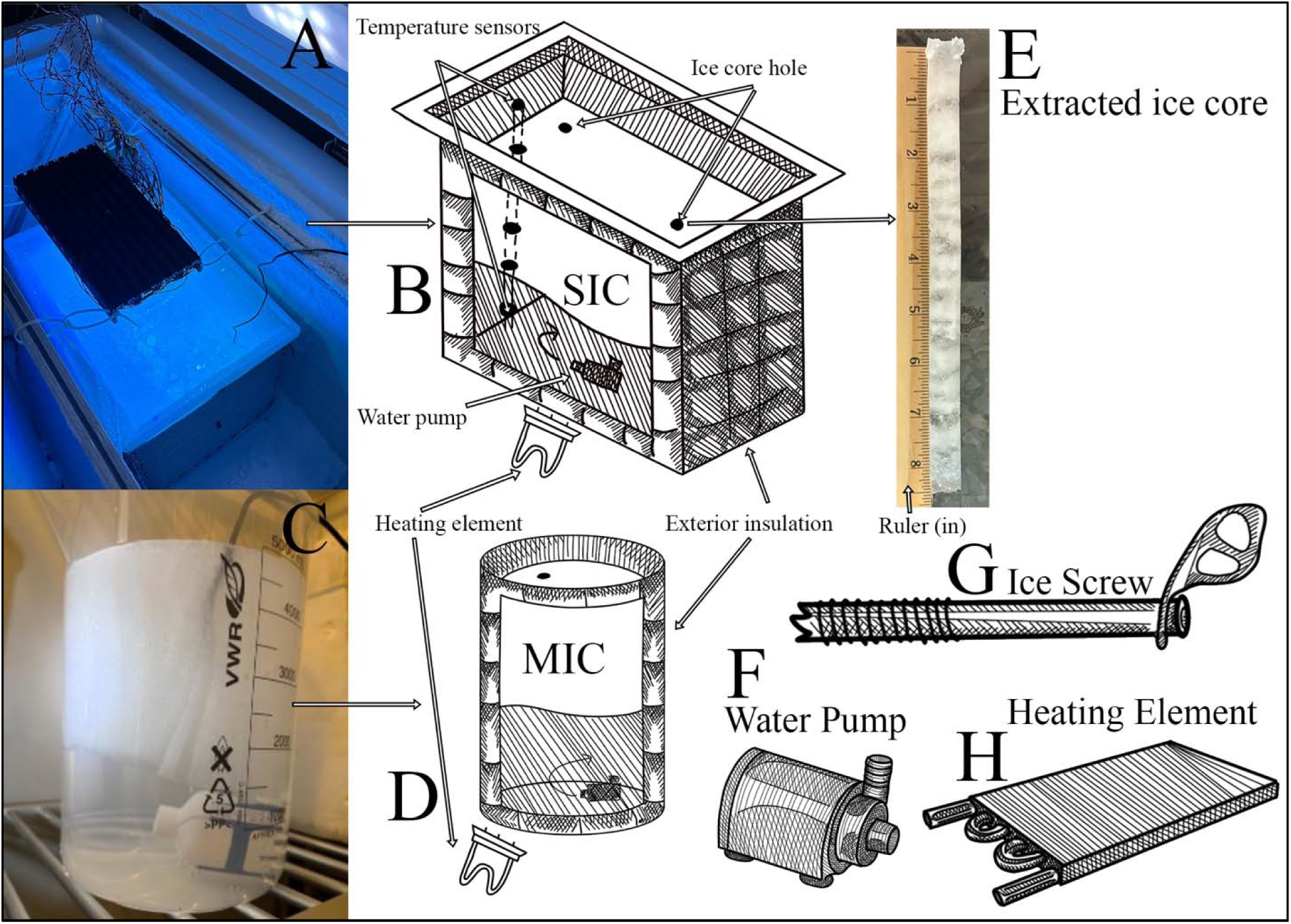
Schematic depicting the design & construction of the Square-Icecosm (SIC) and the Mini-Icecosm (MIC). (A) Picture of the SIC with the ice chest freezer from above. (B) Schematic drawing of SIC construction depicting the exterior insulation, underneath heating element, temperature sensors anchored throughout, interior water pump, and ice core holes. (C) Picture of the MIC extracted from its exterior insulation to observe ice growth and the interior pump. (D) Schematic drawing of the MIC depicting the exterior insulation jacket, interior water pump, and underneath heating element. (E) Example of an ice core extracted from the SIC next to a basic ruler (in). (F) Drawing of a submersible water pump used within the SIC for gentle circulation. (G) Drawing of a basic ice screw used to extract small ice cores from both icecosms. (H) Drawing of the conductive heating element placed underneath the icecosms, connected to an external heat source.

**Table 1:** Temperature & Salinity characteristics of icecosms. Square-Icecosm (SIC) data is reported from each sampling day where possible (A), while Mini-Icecosms (MICs) are separated into 1) mixed cultures, 2) *N. frigida* and 3) *P. glacialis* (B).

| SIC |  |  |  |  |  |  |  |  |  |  |
| --- | --- | --- | --- | --- | --- | --- | --- | --- | --- | --- |
| Day | Ice Depth (cm) | Top (°C) | Mid (°C) | Bot (°C) | Water (°C) | Air (°C) | Top (PSU) | Mid (PSU) | Bot (PSU) | Water (PSU) |
| 0 |  | — | — | — | — | — | — | — | — | — |
| 5 | 11.43 ± 0 | -3.67 | -1.67 | -2.00 | -2.17 | -10.78 | — | — | — | — |
| 7 | 10.2 ± 0 | -5.28 | -2.17 | -2.17 | -2.39 | -13.06 | 39.00 | 22.00 | 25.00 | — |
| 9 | 12.7 ± 1.8 | -6.50 | -3.70 | -2.50 | -2.70 | -11.50 | — | 29.00 | — | 60.00 |
| 14 | 14.6 ± 1.8 | -7.50 | -5.40 | -4.40 | -3.10 | -9.40 | 19.00 | 28.00 | 24.00 | 61.00 |
| 16 | 21 ± 1.8 | -8.00 | -5.70 | -4.80 | -3.20 | -11.40 | 25.00 | 28.00 | 25.00 | — |
| 21 | 22 ± 7 | -8.17 | -6.00 | -5.06 | -3.50 | — | — | — | — | 65.00 |
| 30 | 20.6 ± 1.3 | -8.17 | -5.89 | -5.39 | -3.67 | — | 4.71 | — | 4.31 | 67.00 |

B
| MIC |  |  |  |  |  |  |  |
| --- | --- | --- | --- | --- | --- | --- | --- |
| Core Group | Ice Depth (cm) | Top (°C) | Middle (°C) | Bottom (°C) | Top (PSU) | Middle (PSU) | Bottom (PSU) |
| Mixed | 10.50 | -8.50 | -6.40 | -3.20 | 11.56 | 17.21 | 19.89 |
| *N. frigida* | 12.00 | -8.40 | -6.30 | -3.30 | 14.33 | 10.25 | 18.28 |
| *P. glacialis* | 9.30 | -9.50 | -8.20 | -5.70 | 14.75 | 11.97 | 21.78 |

For the SIC, pre-cooled and sterile artificial seawater (Aquil* medium) with modified nutrients (200 µM NO_3_ ^-^, 200 µM Si, 12.5 µM PO_4_ ^3-^, and L1 metals/ vitamins) adjusted to a salinity of 35 PSU was placed inside a chest freezer to initiate ice formation. For the MICs sterile artificial seawater (ASW; modified from Chen et al. 1996) adjusted to a salinity of 29 was used. Ice formation was monitored continuously. As ice thickened, the temperature of the MIC cooling bath was gradually decreased to promote downward ice growth while avoiding rapid freezing.

### 2.2 Characterization of artificial ice

Physical characterization of the ice and associated biological measurements started in the SIC 5 days after freezing began. Ice cores were extracted using an ice screw (length 22 cm, internal diameter 1.5 cm) and ice depth was measured based on core length (Figure 1). Core length was measured using a ruler and each core was sectioned into a top, middle, and bottom section. Core sections were melted into chilled hypersaline (88 PSU) Aquil* media to reduce osmotic shock and final salinity of the melt water was measured using a refractometer (EXTECH Instruments RF20). The salinity of the ice-core derived melt water was calculated as follows:

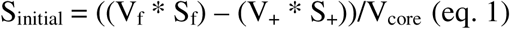

where V_f_ is the final volume of the melted core plus the hypersaline seawater, S_f_ is the total final salinity, V_+_ is the volume of seawater added, S_+_ is the salinity of the added seawater (88 PSU), and V_core_ is the calculated initial core volume (V_f_ - V_+_). Under-ice seawater was sampled through an open ice screw hole using a 10 mL volumetric pipette (VWR 75816-100) and salinity was measured accordingly.

The ice within the MIC was characterized ∼72-hrs after freezing began and ice thickness was measured via visual assessment along the side of the icecosm. The salinity of the melted core sections was measured as conductivity (Mettler Toledo Seven Compact Conductivity S230 with Mettler Toledo In-Lab 731-ISM) directly from the melted core. Under-ice ASW salinity was sampled through an open ice screw hole using tubing attached to a syringe and measured accordingly.

### 2.3 Algae maintenance and cell incorporation into the icecosms

The SIC was inoculated with the polar diatom *F. cylindrus*, selected for its affinity for the sea ice environment and ice-edge blooms (Otte et al. 2023). Prior to inoculation, cultures were grown as sterile, semi-continuous batch cultures, maintained in mid-exponential growth phase in Aquil* medium with modified nutrients (40 µM NO_3_ ^-^, 40 µM Si, 2.5 µM PO_4_ ^3-^ with Aquil* metals/ vitamins) under diel (12:12-hr light:dark) lighting at 80 µmol photons m^-2^ s^-1^ provided by an LED light strip. Prior to inoculation of the tank, tests were conducted to ensure that the impeller pump action had no significant impact on cell growth and cellular integrity (data not shown). The cells were introduced to the icecosm at 4 °C and the freezer temperature was lowered to ∼ -10 °C to initiate ice formation. Ice was allowed to form for 5 days and continued for up to 30 days after. The light panel within the SIC consisted of a 20 cm long and 10 cm wide multi-wavelength panel installed ∼10 cm above the water surface. A wavelength spectrum was selected to mimic available light at 5 m in clear water, and the LED board was controlled by an Arduino (Mega 2560) programmed to form a sinus curve to mimic realistic changes in solar irradiance over a day (Supplementary Figure S1A & B, 12:12-hrs light:dark). Maximum irradiance was 167 µmol photons m^-2^ s^-1^ at peak light and 88.7 µmol photons m^-2^ s^-1^ over the light period.

For the MIC, two diatoms, *Porosira glacialis* (Antarctic, centric) and *Nitzschia frigida* (Arctic, pennate) were grown to high cell densities in ASW F/2 + Si media at 1 °C under diel (12:12-hr light:dark) lighting at a salinity of 29.6. These cells were placed into ASW F/100 + Si media at 0 °C to acclimate to icecosm conditions for 4 days in the dark. These species were selected based on their inherent differences in shape and their occurrence within or near the ice matrix in their natural environment, with the goal of assessing differences in incorporation of presumed suitable (*N. frigida*) and presumed unsuitable (*P. glacialis*) cells into the MIC. *P. glacialis* is reported as a primarily pelagic species, however, is known to occur in close association with sea ice (Pike et al. 2009). Our rationale is that *N. frigida* is known to be strongly associated with the ice environment and blooms regularly within the ice (Michel et al. 2002). Cell survival in the presence of the submerged pump was tested prior to the experiment and exposure over several hours and showed no impact on cell integrity or subsequent population growth (data not shown). Nutrients were added at 1/50th strength of the F/2 + Si nutrients in the ASW (salinity of 29) in the icecosms, as they become concentrated within ice brine channels through the same processes as salts (Arrigo 2014; Granskog et al. 2003; Melnikov et al. 2003). Three individual MICs were inoculated with two MICs containing monoclonal cultures of *P. glacialis* and *N. frigida*, and a third was inoculated with a mix of both diatoms. Diatoms were inoculated into the icecosms in darkness while frazil ice crystals began to form for a final initial icecosm cell density of 500-7000 cells mL^-1^. Ice was allowed to grow for ∼72-hrs, or until ∼10 cm of ice was formed, and then remained stable for ∼24-hrs. Icecosms were sampled after five days after the start of freezing. Please note that MICs were kept in the dark, while in the SIC algae were grown under dynamic irradiance. To control for potential mortality in extended darkness in the MICs, cells from the inoculum were maintained in liquid culture in the light and in the dark in an incubator (Sanyo MLR-351) alongside the MICs. Mortality was assessed using the mortality assay (Samuels et al. 2021) by counting live and dead cells from the under-ice water, the inoculation culture maintained in the light in the incubator, and the inoculation culture maintained in the dark in the incubator (Supplementary Table S2). Cells were stained with 2% Evans Blue stain, as in Samuels et al. (2021), adapted for these species by using a staining time of 1-hr ± 15-min. Samples were concentrated ∼15-fold by centrifugation in an Eppendorf Centrifuge 5424R (8-min, 800 RCF, 2 °C), mounted onto Sedgewick-Rafter cell counters and visualized by standard light microscopy. Live cells presented as pale green or green-gold, while dead cells were stained light grey-blue to dark-blue. Due to large differences in shape, *N. frigida* and *P. glacialis* cells were differentiated visually in the mixed icecosm and cultures.

### 2.4. Distribution & Physiology of *F. cylindrus* within the SIC

Once ice was formed in the SIC, *F. cylindrus* was sampled on day 5, 7, 9, 14, 16, 21, and 30 (Supplementary Table S1). Cores were collected either prior to the photoperiod or at dim light (<16 µmol photons m^-2^ s^-1^). On each day except for day 5 (where a whole, un-sectioned core was collected), ice cores were measured and either split into top, middle, and bottom sections or kept whole, transferred into 50 mL falcon tubes filled with hypersaline seawater, and allowed to melt on ice in the dark (∼4-hrs). To compare melting techniques and the immediate impact on photophysiology (hypersaline seawater addition vs. core melted with no addition), whole un-sectioned cores were measured using Fast Repetition Rate fluorometry (FRRf, LabSTAF, Chelsea Technologies) on day 14. The underlying Aquil* seawater was sampled at the same time as ice core sampling using a 25 mL pipette fed through an open core-hole. *F. cylindrus* distribution within melted cores and under-ice water was measured using a flow cytometer (Beckham Coulter) equipped with a blue laser (488 nm) to excite chlorophyll *a*. Fluorescence emissions were measured using a red (695/50 nm) filter for chlorophyll *a*. Intensity of the single cell fluorescence (Chl *a* cell^-1^), number of cells, size of cells (forward scatter; FWC), and internal complexity (side scatter; SSC) were determined.

Photophysiology of the algae was assessed via FRRf on days 7, 9, 14, and 16 (Supplementary Table S1). FRRf measurements were collected within the same 3-hr sampling window in the early morning. Samples were dark-acclimated during the melting process (up to 4-hrs). Photosynthesis-irradiance (P vs. E) curves were generated by sequential 45-s exposures to increasing irradiance from 0 to 600 µmol photons m^-2^ s^-1^. Maximum irradiance was chosen based on ice core section; cells in the top and middle ice layers were noticeably photo-inhibited at lower irradiances, thus maximum irradiance was lowered compared to the bottom ice layer and the underlying water. Eleven chlorophyll fluorescence yields (ChlF) were obtained for each light step for which the mean of the last five data acquisitions at each light level were used for data assessment (Schuback et al. 2017). Rates of photosynthetic electron transport (JVPII) were calculated in the FRRf RunSTAF software according to (Oxborough 2022):

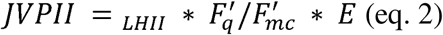

Where ⍰_LHII_ is the absorption coefficient of PSII light harvesting, F_q_’ = F_m_’ - F’, F_mc_’ is the baseline corrected value of Fm’, and E is irradiance (µmol photons m^-2^ s^-1^). Data were further processed in RStudio (R version 4.5.0) and P vs. E curves were fit according to the relationship described by Platt et al. (1980), which accounts for photoinhibition. Modeled values of maximum JVPII and the initial slope of the P vs. E curve (⍰, µmol electrons (µmol photons)^-1^ m^-1^) were used to calculate the light saturation parameter, E_k_ (JVPII_MAX_ / ⍰; µmol photons m^-2^ s^-1^). Values of JVPII_MAX_ were then normalized to cell concentration to obtain rates of electron transport per cell (ETR_cell_) across core sections.

The Normalized Stern-Volmer Non-Photochemical Quenching (NPQ_NSV_) coefficient in the dark was calculated as NPQ_NSV_ = (F_m_’/F_v_’) -1 = F_0_’/F_v_’. The rate of Q_a_ reoxidation, 1/τ, was measured during the relaxation step of the dark-adapted single-turnover measurement described in Oxborough 2022. See Table 2 for a list of other physiological terms discussed.

**Table 2:** Physiological terms used throughout this manuscript.

| Parameter | Definition | Calculation | Unit |
| --- | --- | --- | --- |
| $F(')$ | Any ChlF point between $F_o$ and $F_m$ | | unitless |
| $F_m(')$ | Maximum fluorescence in a dark-acclimated sample (light acclimated sample) | | unitless |
| $F_o(')$ | Base fluoresce in a dark-acclimated sample (light acclimated sample) | | unitless |
| $F_v(')$ | Variable fluorescence in a dark-acclimated sample (light acclimated sample) | $F_m(') - F_o(')$ | unitless |
| $F_v/F_m$ | Quantum photochemical efficiency of PSII | $F_m - F_o / F_m$ | unitless |
| $NPQ_{NSV}$ | Normalized Stern-Volmer Non-Photochemical Quenching | $(F_m' / F_v') - 1 = F_o' / F_v'$ | unitless |
| $\sigma_{PSII}$ | Functional absorption cross section | | $\text{nm}^2$ |
| $1/\tau$ | Rate of reoxidation of $Q_a$ | | $\mu\text{s}^{-1}$ |
| $\Phi$ | Photosynthetic efficiency under light limitation | Derived from P vs E curves, Fit from Platt et al. 1980 | |
| $E_k$ | Light-saturation parameter | $JVP_{II}m / \alpha$ | $\mu\text{mol photons m}^{-2} \text{ s}^{-1}$ |
| $JVP_{II}$ | Maximum PSII flux per unit volume | Fit from Platt et al., 1980 | $\mu\text{mol electrons m}^{-3} \text{ d}^{-1}$ |
| $ETR_{\text{cell}}$ | Electron transport rate (normalized to cell concentration) | | $\text{nmol electron cell}^{-1} \text{ d}^{-1}$ |

### 2.5. Characterization of diatoms in the MIC

All sampling in the MICs was done after 4 days in the dark and frozen in ice. Separate cores were taken for diatom presence, chlorophyll quantification, and growth assays. To measure diatom presence in ice cores and under-ice water, ice cores were taken with an ice screw (Black Diamond Express; length 19 cm, internal diameter 1.4 cm). The ice screw was rinsed with 70% ethanol between each use. Ice cores were measured and either split into a top and bottom section or kept whole and allowed to melt at 1 °C to measure diatom presence and growth. Cells in the underlying ASW were sampled at the same time using tubing attached to a syringe from an open core-hole. Once cores were fully melted (∼4-hrs), cells were counted via microscopy using a Sedgewick-Rafter cell counter.

For chlorophyll measurements, full or partial ice cores were melted in the dark at 1 °C. Cells were filtered on to GF/F filters, and frozen in foil at -20 °C. To extract chlorophyll, filters were placed in 15 mL sterile falcon tubes alongside four 5 mm steel beads and 10 mL 96% ethanol, shaken for 5-mins to disrupt the filter and subsequently incubated overnight in the dark at 4 °C. After extraction, tubes were centrifuged at 3500 rpm for 10-mins to pellet debris. Fluorescence from chlorophyll extracted into the supernatant was measured at excitation and emission wavelengths of 433 and 673 nm, respectively, using a spectrophotometer (**Perkin** Elmer Luminescence Spectrometer LS 30 chlorophyll *a* concentration was then calculated using an internal calibration curve (0.1182x + 0.0984).

To assess the potential for pelagic growth following freezing into the ice and being thawed, cores were taken and used to inoculate fresh F/2 media at 2 °C under diel (12:12-hr light:dark) lighting at a salinity of 29.6 (identical to initial culturing conditions). At least one core was kept intact; others were sectioned into top and bottom layers. These cultures were reintroduced to light (30 µmol photons m^-2^ s^-1^) and allowed to grow for up to 57 days. After 7, 20, and 57 days, cells were stained with Lugol’s dye (Hällfors 1979) and counted via light microscopy using a Sedgewick-Rafter cell counter and growth rates were determined using an exponential fit function.

## 3. Results

### 3.1. Physical icecosm characteristics

Ice in the SIC grew progressively throughout the duration of the experiment and ranged from 11.43 cm (n = 1) after 5 days of freezing to a maximum of 21 ± 1.8 cm (n = 2) after 16 days of freezing (Supplementary Table S1). Vertical temperatures in the SIC ranged from ∼ -8 °C in the upper ∼2 cm to ∼ -2.5 °C in the bottom ∼2 cm (Table 1A). Direct measurements of pure brine salinity were largely unsuccessful due to rapid ice-core melting immediately post-extraction, which rendered brine leakage indistinguishable from the melting ice matrix. Please note that the salinity used for the photophysiological measurement was higher due to the addition of hypersaline seawater, as indicated in the methods and was on average 43 ± 7 PSU. To see the theoretical calculated bulk melted salinity without the addition, please refer to Table 1. Salinity in the underlying water column increased to 67 PSU on day 30. Subtle structural banding was observed in the SIC cores, potentially attributable to the freeze/thaw cycles of the chest freezer (Figure 1E).

Three MICs were operated simultaneously. Total ice thickness varied among the chambers, ranging from 9.3 cm in the *N. frigida* MIC to 12.0 cm in the *P. glacialis* MIC (Table 1B). A vertical temperature gradient was observed within the mixed species MIC and the *N. frigida* MIC, ranging from ∼ -8.5 °C in the upper ∼2 cm to ∼ -3.2 °C in the bottom ∼2 cm. The *P. glacialis* MIC exhibited a similar but slightly colder gradient spanning from -9.5 °C in the upper ice to -5.7 °C in the bottom ice. Bulk salinity of the melted ice cores ranged from 13.8 to 19.6 in the MIC’s, which is a reduction from the starting ASW salinity of 29. Salinity in the underlying water reached up to 54 PSU.

### 3.2. Cell distribution and optical properties: SIC

Cellular concentrations, optical properties, and photophysiological traits (electron transport and quenching parameters) were monitored across vertical ice-core gradients and in the underlying water over 30 days. Due to limited sampling replication (n = 1), data are presented descriptively without statistical testing. Following the initial inoculation into the SIC for a final concentration of ∼380,000 (± 17,000) cells mL^-1^ (n = 2), *F. cylindrus* immediately incorporated into the ice column during ice formation, as confirmed by a whole-core extraction on day 5 (data not shown). Distinct vertical stratification was observed in our first sectioned ice-core on day 7 with the highest cell densities concentrated in the bottom ice layer (Figure 2A). In this bottom section, cell densities increased steadily from ∼ 170,000 ± 1,600 cells mL^-1^ on day 7 to ∼ 240,000 ± 5,300 cells mL^-1^ on day 14. Changes in cell density in the middle ice layer was more moderate, increasing from ∼115,000 ± 2,700 cells mL^-1^ on day 7 (187,000 ± 5,700 cells mL^-1^) on day 9 before leveling off at by day 14 (172,000 ± 3,400 cells mL^-1^). In contrast, cell counts in the top section remained largely stagnant (35,950 – 81,278 cells mL^-1^) or declined throughout the duration of the experiment (Figure 2A). Between days 14 and day 16, a general decline in cell abundance was observed across all ice-sections, with densities remaining relatively constant thereafter until the end of the experiment on day 30. By day 16, 41.8% of the main *F. cylindrus* population was concentrated in the bottom ∼2 cm of ice, 29.3% in the middle ∼2 cm, 14.4% in the top ∼2 cm, and 14.5% of cells remained in the underlying water. This distribution shifted slightly by the end of the experiment (day 30), resulting in a final distribution of 37% of cells contained in the bottom layer, 36.8% of cells in the middle layer, 23.4% of cells within the top layer, and 2.8% of cells in the underlying water.

**Figure 2:**
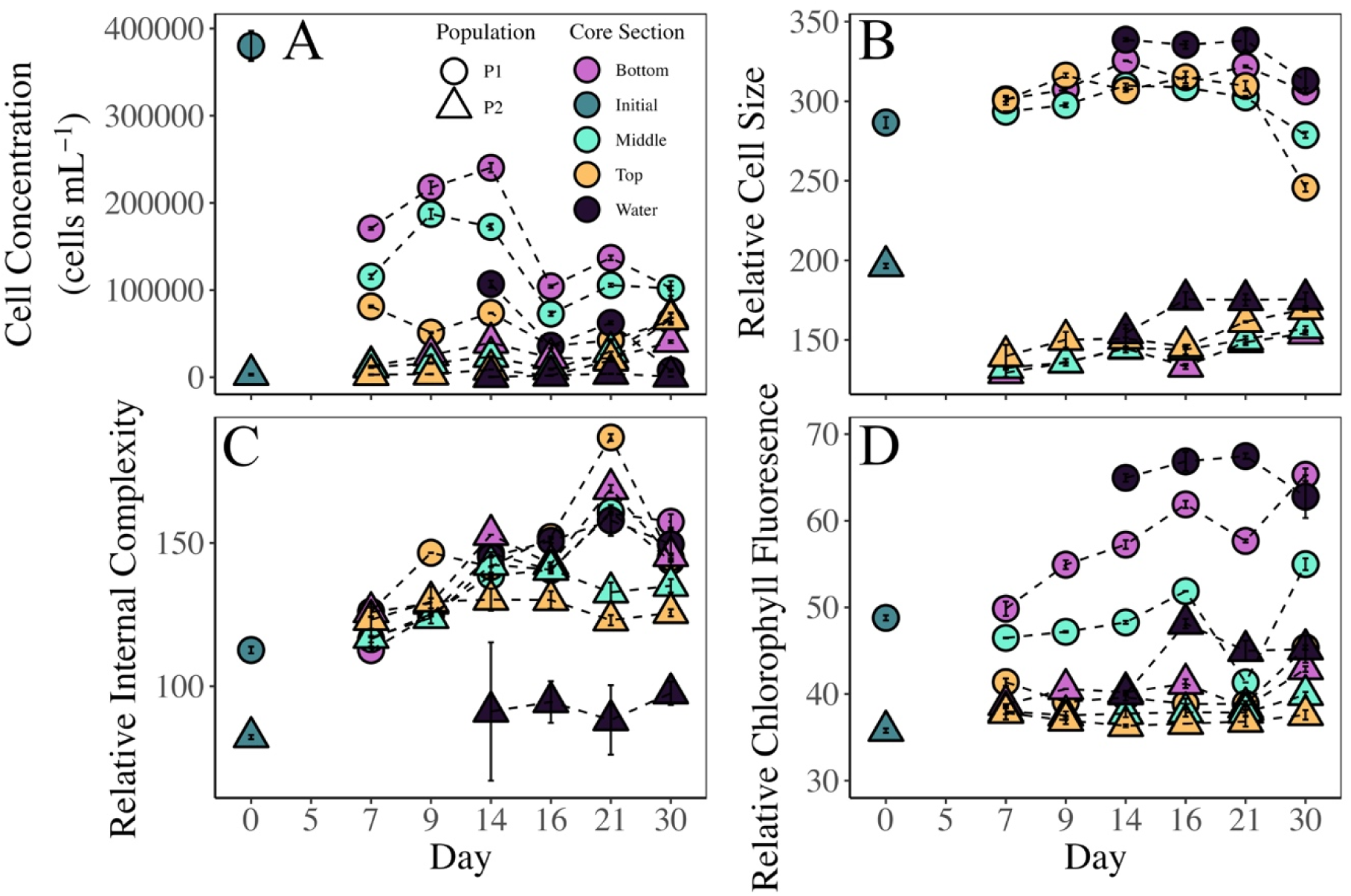
Flow cytometric cell concentration and optical properties of *F. cylindrus* within the SIC over 30 days. (A) Cell concentration in the initial inoculum (teal), underlying water (black), bottom ice layer (purple), middle ice layer (light blue), and top ice layer (yellow). The main population of *F. cylindrus* is denoted as circles and the smaller, sub-population is denoted as triangles. (B) Relative cell size (FSC); (C) relative internal complexity (SSC); and (D) relative chlorophyll *a* fluorescence.

A major biological outcome of cell entrapment into the ice was the emergence of a morphologically distinct sub-population, designated as P2, which was absent from both the underlying water column and the initial inoculum (Figure 2A). Flow cytometric analysis revealed that the P2 sub-population exhibited an approximate 50% reduction in relative cell size compared to the original population, characterized by a distinct shift in forward scatter (FSC, Figure 2B). While this sub-population was initially absent or in very low abundance on day 5 (0.6% of the total cell population), it increased in concentration within the ice over the 31-day incubation period. On average, this smaller sub-population was distributed across all ice sections, though it became increasingly prominent across the ice layers as the experiment progressed. The spatial distribution of the sub-population was heavily biased toward the ice environment; by day 30, the sub-population accounted for 9%, 15.1%, and 14.7% of the total cell population in the bottom, middle, and top ice sections, respectively, while only accounting for 0.2% of the cell population in the underlying water. The increase in concentration of the sub-population within the underlying water during the incubation period may be attributed to brine leakage during core sampling.

The optical properties of the *F. cylindrus* cells shifted significantly over the duration of the experiment as they acclimated to the environmental gradients within the SIC (Figure 2B-D). Relative cell size (FSC) of the main *F. cylindrus* population remained stable for the first two weeks but showed a marked decrease by the final day of sampling, with the largest reduction occurring in the top section of the ice (Figure 2B). Simultaneously, the relative internal complexity (SSC) of both the P1 and P2 populations increased throughout the course of the experiment, indicating changes in cellular density or granularity (Figure 2C). Per-cell chlorophyll *a* fluorescence also showed an increase in the bottom and middle layers of the ice over time, reaching peak values by day 30 that were approximately 1.2 times higher than the measurements taken on day 7 (Figure 2D).

### 3.3. Photophysiology: SIC

The photophysiological response of *F. cylindrus* was strongly influenced by both vertical positioning within the ice matrix and the duration of entrapment (Figure 3A-D, sampled days 7 through 16). The maximum quantum yield of PSII (F_v_/F_m_) was consistently highest in the bottom ice layer, though it exhibited a steady decline of 31% from 0.32 on day 7 to 0.22 by day 16 (Figure 3A). This downward trend was more pronounced in the middle layer where F_v_/F_m_ decreased by 38% (from 0.21 to 0.13), and most severe in the top layer with a 77% reduction from an initial low of 0.13 to a near-baseline value of 0.03 by day 14. In contrast, cells in the underlying water column maintained relatively stable photosynthetic health, with only slightly decreasing F_v_/F_m_ by 7% (from 0.42 to 0.39) between inoculation and day 16. As photosynthetic efficiency declined within the ice matrix, energy dissipation through quenching processes increased significantly. The highest values of normalized Stern-Volmer non-photochemical quenching (NPQ_NSV_) were recorded in the top ice layer, ranging from 5.3 to 36.2 (Figure 3B). The functional absorption cross-section of PSII (area of light absorption, σPSII (nm^2^ PSII^-1^)), remained relatively stable across ice sections and over time (Figure 3C). Rates of Q_a_ reoxidation, (1/τ, ns), were variable and showed no consistent trend throughout the incubation (Figure 3D). 1/τ values were overall lower in the top layer of the ice on days 7 and 9, however after day 9, 1/τ was unable to be calculated due to high signal noise as fluoresce induction towards F_m_’ nearly fails.

**Figure 3:**
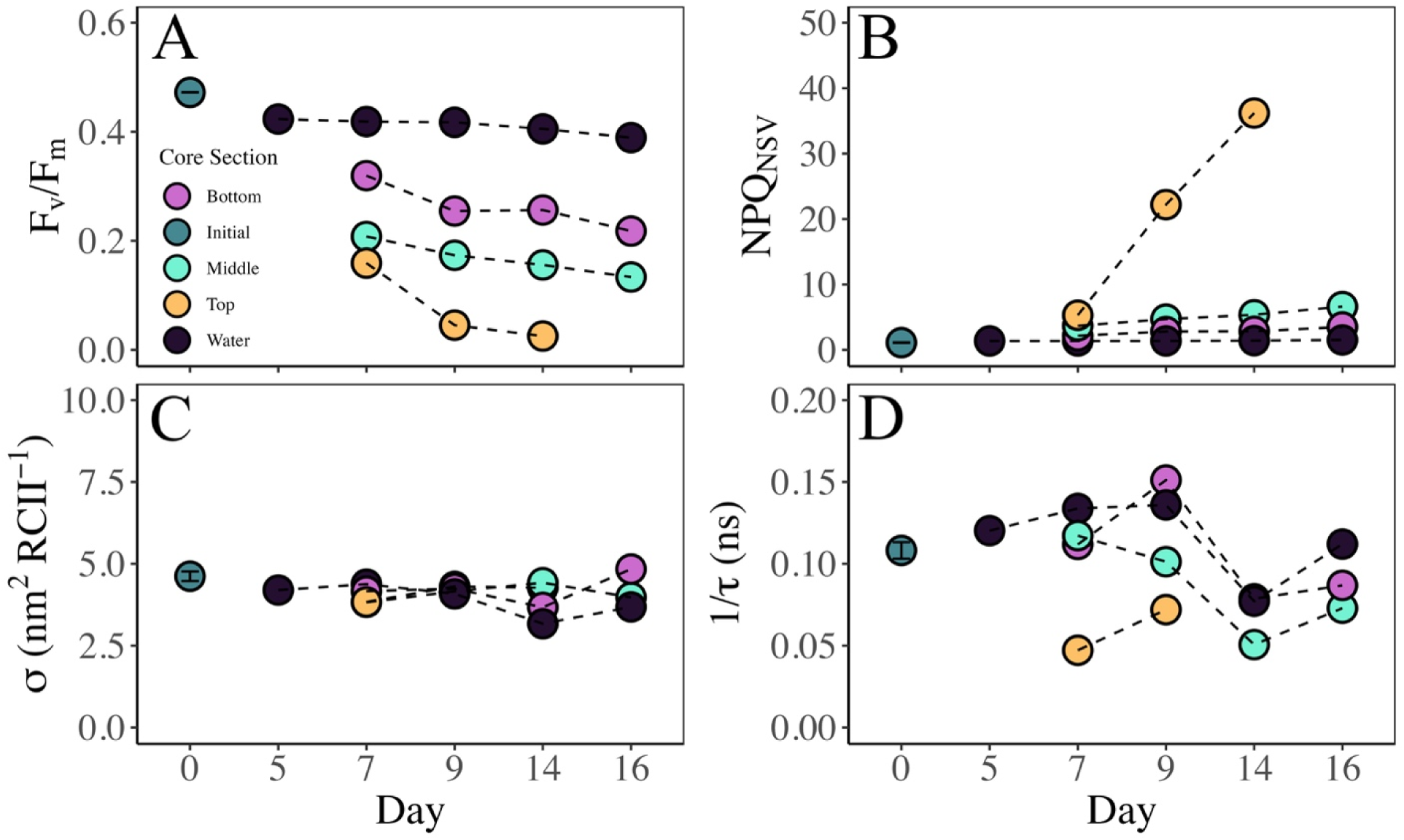
Photophysiologcal parameters of *F. cylindrus* within the SIC over 16 days. (A) Quantum photochemical efficiency (F_v_/F_m_) in the initial inoculum (teal), underlying water (black), bottom ice layer (purple), middle ice layer (light blue), and top ice layer (yellow). (B) Normalized Stern-Volmer Non-Photochemical Quenching (NPQ_NSV_). (C) Functional absorption cross sectional area of PSII photochemistry (sigma, nm^2^ PSII^-1^). (D) Rate of reoxidation of the primary electron acceptor, Q_a_ (ns).

Photosynthetic kinetics, derived from FRRf light-response curves (P vs. E) curves, further highlighted the physiological changes of algae entrapped within the different ice layers. The initial slope, ⍰ (µmol electrons µmol photons^-1^ m^-1^; representing low-light photosynthetic efficiency) remained consistently higher in the bottom layer, however, was overall variable across the core sections (Figure 4A). Excluding data from the unreliable top ice layer, the light saturation parameter, E_k_ (µmol photons m^-2^ s^-1^), was slightly higher in the middle ice layer than in the bottom ice layer and increased in the underlying water (Figure 4B).

**Figure 4:**
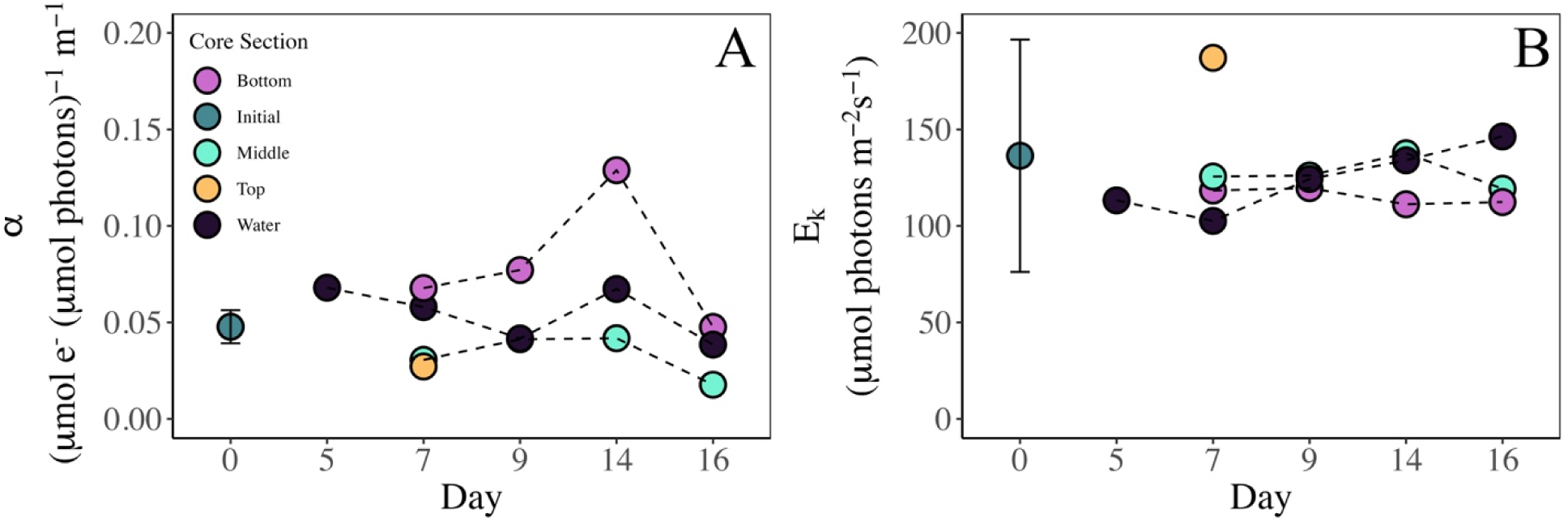
Light acclimation parameters from FRRf for *F. cylindrus* in the SIC over 16 days. (A) Low light photosynthetic efficiency (alpha, µmol electrons µmol photons^-1^ m^-1^) in the inoculum (teal), underlying water (black), bottom ice layer (purple), middle ice layer (light blue), and top ice layer (yellow). (B) Light saturation parameter (Ek, µmol photons m^-2^ s^-1^). FRRf JVPII vs E curves were fit using the Platt et al. (1980) productivity model and Ek was calculated as JVPII_MAX_ /⍰.

Maximum rates of electron transport per cell (ETR_cell_) remained relatively stable within the middle (ranging from 0.026 to 0.03 nmol electrons cell^-1^ d^-1^) and bottom ice layers (ranging from 0.04 to 0.05 nmol electrons cell ^-1^ d^-1^) throughout the incubation (Figure 5). However, in the underlying water, ETR_cell_ fluctuated, initially decreasing until day 9 before increasing by day 16. In the upper ice column, the maximum rate of electron transport could not be reliably determined. Although cells were physically present in this top section, the signal-to-noise ratio was insufficient to yield valid physiological data, highlighting the high stress these cells experience in the upper ice column.

**Figure 5:**
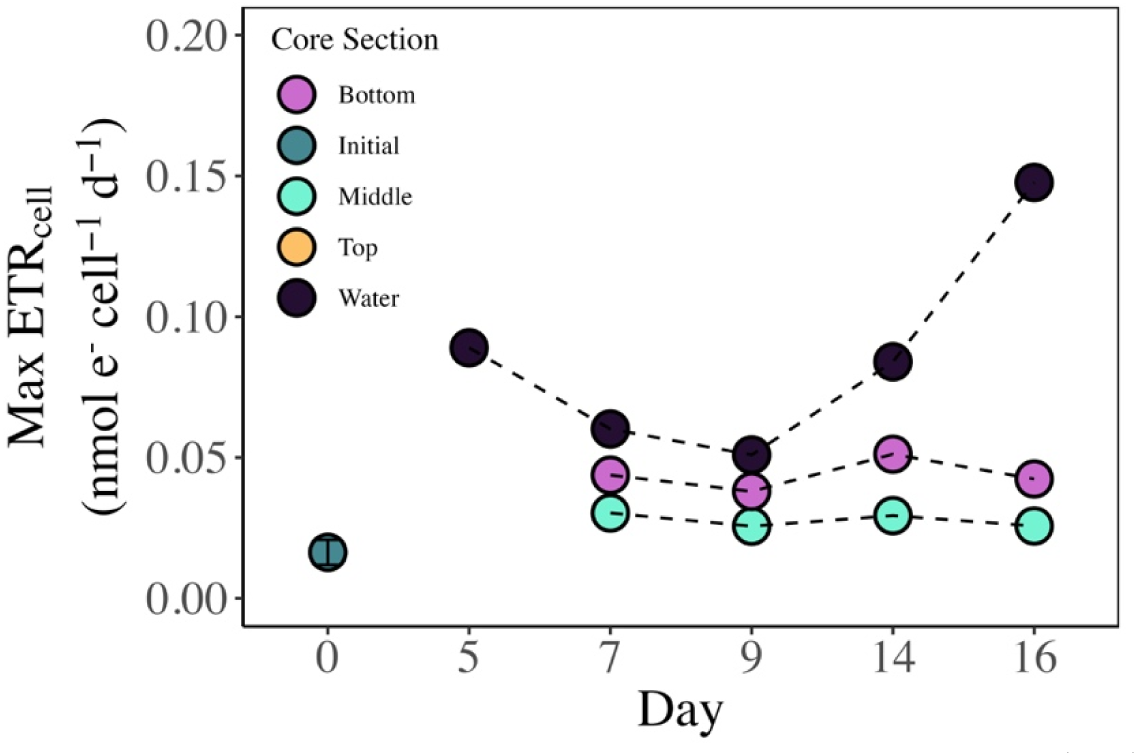
Maximum rates of electron transfer (ETR_cell_, nmol electron cell^-1^ d^-1^) in the inoculum (teal), underlying water (black), bottom ice layer (purple), middle ice layer (light blue), and top ice layer (yellow) from the SIC. FRRf P vs E curves were fit according to the Platt et al., (1980) productivity model and maximum rates of JVPII were normalized to cell concentration within the ice.

To evaluate potential measurement artifacts associated with ice-core processing methods, two additional whole (un-sectioned) ice cores were collected on day 14 to compare melting protocols: one core was melted with the addition of buffered 88 PSU seawater to mitigate osmotic shock, while the other was melted directly without seawater addition (Figure 6A-G). No substantial differences were observed in F_v_/F_m_ (0.21 vs 0.22), NPQ_NSV_ (3.7 vs 3.5), or σPSII (4.5 vs 4.3 nm^2^ PSII^-1^). However, both 1/τ and ⍰ were depressed under the direct freshwater (“FW”) melt condition, whereas E_k_ and maximum ETR_cell_ were enhanced. These specific kinetic divergence patterns indicate that localized reductions in salinity during processing may include rapid osmotic stress, artificially altering photosynthetic kinetics (⍰ and E_k_) to resemble a high-light acclimated state.

**Figure 6:**
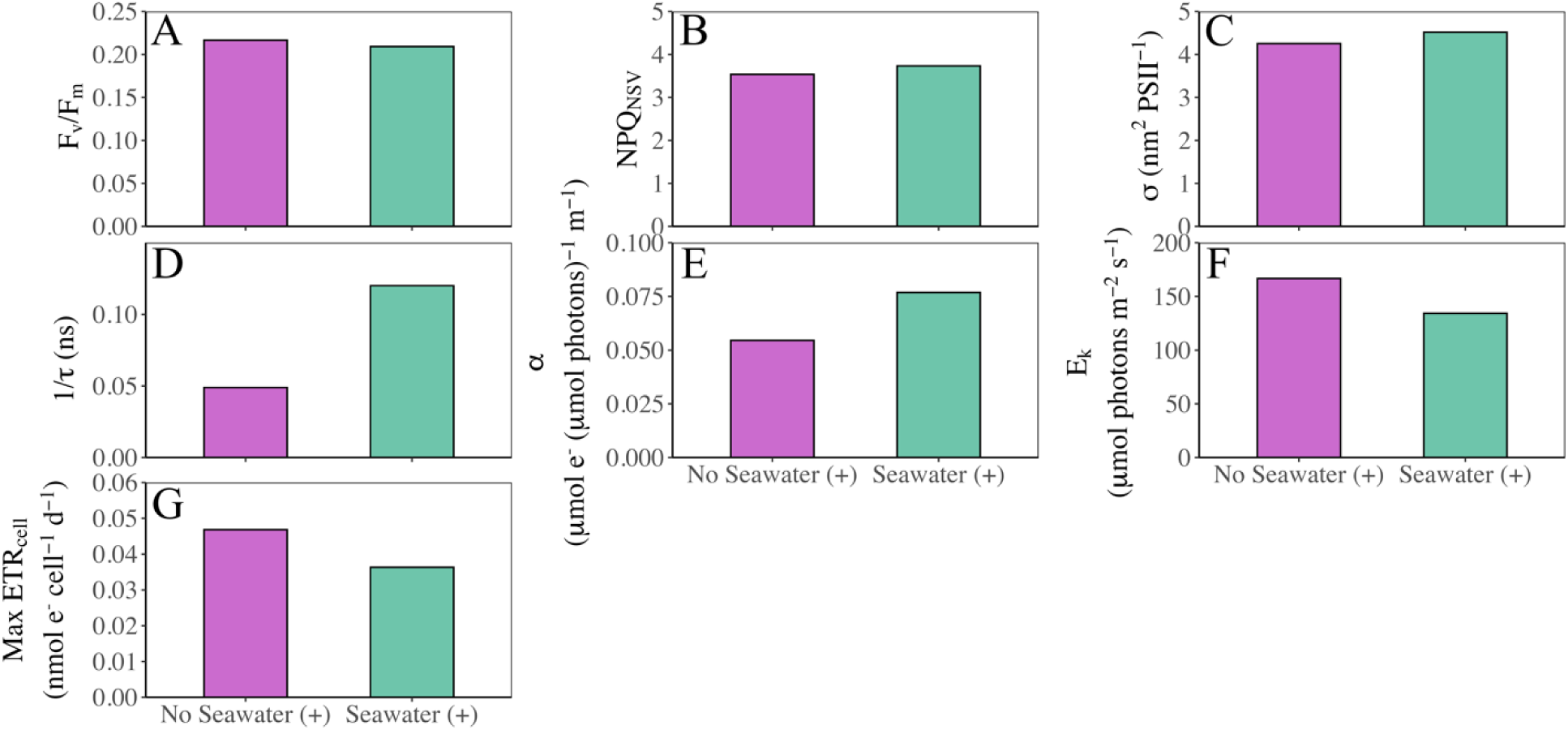
Physiological comparison of SIC ice core melting techniques. Cores melted in hypersaline (88 PSU) Aquil* seawater are represented in turquoise, and cores melted without any seawater addition are represented in purple. (A) Quantum photochemical efficiency (F_v_/F_m_). (B) Normalized Sten-Volmer Non-Photochemical Quenching (NPQ_NSV_). (C) Functional absorption cross sectional area of PSII photochemistry (σPSII). (D) Rate of reoxidation of the primary electron acceptor, Q_a_ (1/τ, ns). (E) Slope of the light-limited region of the FRRf-derived JVPII vs E curve (⍰, µmol electrons µmol photons^-1^ m^-1^). (F) Light saturation parameter (E_k_, µmol photons m^-2^ s^-1^). (G) Maximum rate of electron transport (ETR) normalized to cell concentration (nmol electron cell^-1^ d^-1^).

### 3.4. Diatom incorporation into ice & melt survival: MIC

Successful incorporation of both *P*. *glacialis* and *N*. *frigida* into the ice matrix was confirmed by chlorophyll *a* measurements (Figure 7). Both diatom cultures used to inoculate the MICs contained few dead cells (1-8%, Supplementary Table S2). Due to biomass limitations, cell quantification could not be performed alongside chlorophyll *a* (Chl *a*) extractions on the MIC ice cores; however, Chl *a* concentration served as a robust proxy for cellular integration into the ice.

**Figure 7:**
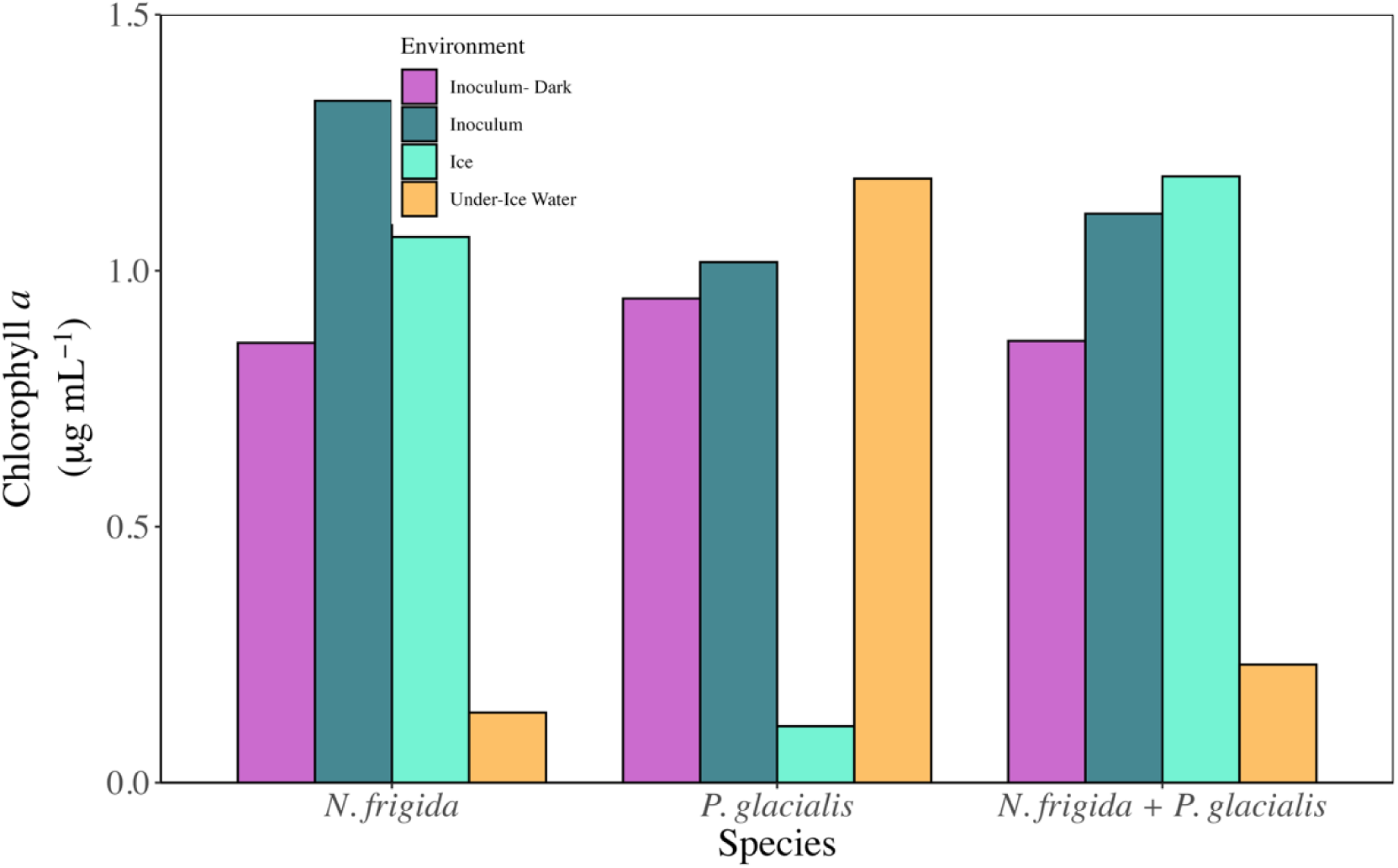
Chlorophyll *a* (Chl *a*) concentration obtained from monoculture MICs with *N. frigida* (left), *P. glacialis* (middle), and the mixed culture MIC with both species (right). Chl *a* from the inoculum kept in the dark alongside the MIC incubations is represented in purple. Chl *a* from the inoculum cultures maintained in the light during the MIC incubation is represented by teal. Chl *a* from MIC ice cores is represented by light blue. Chl *a* from the underlying water in the MICs is represented by yellow.

Immediately post-inoculation and prior to freezing, initial Chl *a* concentration in the ASW were 1.33 µg mL^-1^ and 1.02 µg mL^-1^ for *N*. *frigida* and *P*. *glacialis* icecosms, respectively (Figure 7). At the end of the experiment, ice cores had a Chl *a* concentration of 1.06 µg mL^-1^ and 0.11 µg mL^-1^ for *N*. *frigida* and *P*. *glacialis*, respectively. This discrepancy was mirrored in the underlying water column, where post-freezing Chl *a* concentrations were 0.14 µg mL^-1^ and 1.18 µg mL^-1^ for *N*. *frigida* and *P*. *glacialis* icecosms, respectively. In the mixed species MIC, similar trends to the *N. frigida* MIC were observed with 1.18 µg mL^-1^ Chl *a* incorporated into ice and 0.23 µg mL^-1^ in the under-ice water. This is consistent with high incorporation of *N. frigida* into the ice, with few cells remaining in the under-ice water, and poor incorporation of *P. glacialis* into the ice, with most cells remaining in the under-ice water. Chl *a* concentration in the dark-maintained control inoculums decreased by 36%, 7%, and 22% in the *N. frigida*, *P*. *glacialis*, and mixed MIC, respectively, relative to initial baseline value.

To assess mortality within the underlying water column, the proportion of non-viable cells in the under-ice water was compared to control cultures maintained in the dark over the same experimental duration (Supplementary Table S2). Cells maintained in the dark in liquid media at 1 °C showed low mortality over the same time period as the MIC experiment (1-9%). In contrast, mortality in the under-ice water in the MICs was 36-78%. This divergence is consistent with elevated mortality observed beneath the ice being driven by the extreme environmental conditions of the under-ice brine matrix rather than dark stress or culture age. While cellular abundance within the ice matrix was insufficient to directly quantify within-ice mortality rates using cell counts on the volumes sampled in single ice cores, the persistence of Chl *a*, coupled with successful post-melt regrowth, indicates that a viable proportion of the population was successfully incorporated into the ice.

Post-melt population growth dynamics for *N*. *frigida* and *P*. *glacialis* are detailed in Figure 8. All cultures inoculated from a melted ice core demonstrated positive growth with two exemptions: the *N*. *frigida* population collected from the under-ice water column and the *P*. *glacialis* population isolated from the upper half of the ice core. Although a few live cells were detected with Evan’s blue staining in the under-ice water from the *N*. *frigida* icecosm immediately following the experiment (Supplementary Table S1), these cells failed to divide under the culture conditions of this experiment. Conversely, *P*. *glacialis* cells recovered from the under-ice water exhibited a depressed growth rate (0.04 d^-1^), potentially indicating a prolonged lag phase following hypersaline brine exposure or a low initial proportion of dividing cells within the recovered pool. It is also possible that because more *P. glacialis* than *N. frigida* cells were present in the under-ice water, that the larger absolute number of viable *P. glacialis* cells was sufficient to sample live cells, while the smaller absolute number of *N. frigida* cells did not contain enough viable cells due to sampling error – more replication or experiments with higher cell densities could resolve this. Both *N*. *frigida* and *P*. *glacialis* successfully grew post-melt from the mixed culture MIC. Among all cultures obtained from ice cores, *N*. *frigida* growth rates ranged between 0.03-0.06, while *P*. *glacialis* growth rates ranged from 0.04-0.12. Qualitative observations during live-dead cell counts suggest that smaller *P. glacialis* cells experienced higher mortality while larger cells exhibited superior survival; however, high-resolution cell size metrics were insufficient to statistically validate this size-dependent survival hypothesis.

**Figure 8:**
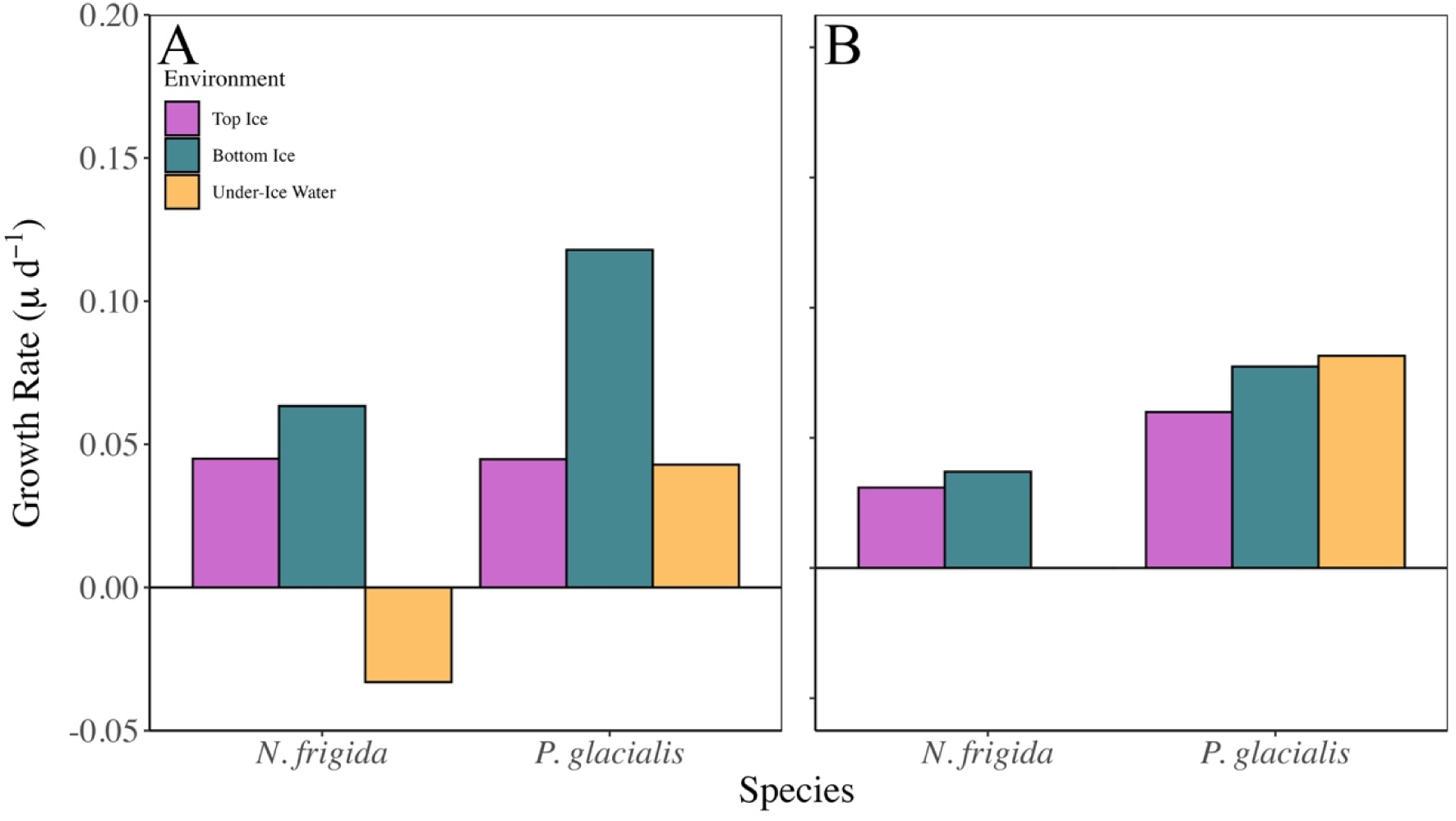
Growth rates (µ d^-1^) of *N. frigida* and *P. glacialis* obtained from the monoculture MIC’s (A) and the mixed culture MIC (B). Purple represents growth rates of cultures inoculated from the top section of MIC ice. Teal represents growth rates of cultures inoculated from the bottom section of MIC ice. Yellow represents growth rates of cultures inoculated from the underlying icecosm water.

## 4. Discussion

The successful development and implementation of the relatively simple and easy-to-build icecosm systems presented here is a significant advancement in our capacity to empirically test the resource economics, environmental limits, and survival abilities of sea-ice algae under controlled conditions. By providing a reproducible, low-cost complement to logistically demanding polar field sampling and expensive, non-reproducible large-scale ice tanks, these microcosms offer an accessible platform for exploring microalgal physiology, ecology, and evolutionary processes in extreme environments. While this framework builds upon previous work based on Yoshida et al. (2020, 2021) and Kennedy et al. (2012), it provides new insights into the physiological and ecological processes of ice-associated diatoms. Our findings demonstrate that these systems yield robust, high-resolution data that capture both in-situ photophysiological acclimation and the subsequent “grow-out” potential of organisms following the critical ice-to-water transition. Although this pilot study represents an initial step in characterizing detailed resource-use efficiencies, the platform establishes a standardized framework for repeatable small-scale investigations into microbial life-history strategies at the physical boundaries of life, aligning with Raven’s broader interest of algae across the globe and into the realm of extraplanetary research where no algae has gone before.

Despite the limited replication in this pilot study, the distinct physiological and distributional responses observed among *F*. *cylindrus*, *N*. *frigida*, and *P*. *glacialis* are consistent with their recognized ecological niches. Notably, *F*. *cylindrus* and *N*. *frigida,* species typically associated with sea-ice habitats (Michel et al. 2002; Otte et al. 2023), reached higher abundances within the ice matrix than *P*. *glacialis*. While *P. glacialis* is considered an ice-associated species, it typically blooms pelagically along the ice edge and occurs within consolidated sea ice only in low concentrations (Pike et al. 2009) (Figures 2A & 7). These distributional differences indicate that the icecosm platform can mimic key habitat preferences and potentially be used in the future to dissect the selection pressures associated with specific aspects of complex environments that underly these preferences. Furthermore, the melt-out experiments confirmed that short term entrapment within the ice does not preclude viability, and highlights that the specific vertical position of entrapment, and the associated temperature and salinity stressors, can exert control on the initial growth rates following the ice-to-water transition. Overall, the icecosms open the possibility of systematically examining how each of these factors affect pelagic growth following melt-out.

In addition to results associated with incorporation into, and release from ice, our study revealed dynamics that occurred within ice-bound populations. The appearance of a morphologically distinct, smaller sub-population of *F*. *cylindrus* (designated as P2), almost exclusively within the ice matrix, indicates an active, acclimatory and selective cell growth response. While *F. cylindrus* is known to adjust osmolyte synthesis and protein turnover to manage low temperature and high salinity (Krell et al. 2007; Uhlig 2011), the formation of a distinct, small-cell morphotype or resting stages within sea ice has not been previously documented to our knowledge. Under stress or prolonged darkness (such as the extended polar night), *F. cylindrus* is known to shift to a metabolic standby state, downregulating cellular processes until conditions become favorable for continued growth (Joli et al. 2024; Morin et al. 2020). However, given that our SIC provided sufficient light and nutrients within the brine network, sustained cell division was maintained. Under the extreme osmotic pressures of the brine channels and or the size restrictions within brine channels, may have supported daughter cells that were significantly smaller than the parent cells initially incorporated from the inoculum. Miniaturization reduces the cellular radius, effectively maximizing the surface-area-to-volume ratio. Biophysically, an increased surface-area-to-volume ratio optimizes the efficiency of transmembrane ion pumps and nutrient transporters, providing a distinct energetic advantage when operating under high brine viscosities and intense osmotic gradients (Irwin et al. 2006; Pan et al. 2024). Further investigation into the characteristics of this sub-population and its growth potential post-melt is required.

Natural sea ice is characterized by steep physiochemical gradients, where brine salinity and temperature within the porous channels are governed by atmospheric and underlying seawater boundaries (Kennedy et al. 2012; Tucker et al. 1992). *In-situ* brine salinity exceeding 200 PSU and temperatures dropping below -20 °C are frequently observed in upper ice layers (Arrigo et al. 2010; Petrich and Eicken 2017), creating microenvironments that are largely hostile to sustained biological activity, despite the capacity of some polar diatoms to maintain minimal photosynthetic function down to -10 °C (Ralph et al. 2005; Thomas and Dieckmann 2002). The interplay between temperature and salinity is a primary driver in structuring the spatial distribution and biological activity through the ice (Kennedy et al. 2012). This relationship is clearly reflected in our physiological profiles from *F*. *cylindrus* in the SIC (Figure 3A). Consistent with observations by Kennedy et al. (2012), maximum quantum yield (F_v_/F_m_) declined towards the top of the ice column where temperatures were coldest and salinities highest. Additionally, the increase in cellular chlorophyll in the lower sections of the ice matrix (Figure 2D) represents a classic photoacclimatory response to low decreasing irradiance, demonstrating that these cells retained functional, responsive metabolic machinery despite low temperatures. Bottom ice layers typically maintain temperature and salinity akin to those of the underlying water column, providing a more hospitable habitat to ice algae (Arrigo 2014), explaining the high concentration of *F. cylindrus* and elevated chlorophyll of *N. frigida* in the bottom ice layer of our icecosms. Notably, this preferential accumulation occurred despite an elevated bulk salinity in the bottom layer, an artifact of our closed system, which prevented the gravity-driven drainage of expelled brine out of the lower matrix, a process that occurs in natural open systems.

Evaluating the system through the lens of a thermodynamic and redox balance can resolve the underlying mechanisms driving the vertical shifts in physiology. Specifically, how light energy is managed when the input (light intensity) outpaces the downstream processing capacity of the enzymatic machinery due to severe environmental bottlenecks. The functional absorption cross-section (σPSII), which represents the functional “antenna” size of the reaction centers of the cells, remained remarkably stable across the ice column (Fig 3C). This structural stability indicates that the physical capacity to capture photons is preserved across the ice-column, even in high-stress zone of the upper ice layers. However, the subsequent conversion and downstream transport of that captured energy was heavily affected by the localized physicochemical environment (Raven and Geider 1988).

In the upper ice layers, where irradiance is highest and self-shading is negligible, the unattenuated energy capture encountered a profound kinetic bottleneck. The low temperature and high salinity crippled downstream enzymatic dark reactions, resulting in a severe “back-pressure” of electrons within the photosynthetic transport chain. When rates of light absorption exceed the processing capacity of the enzymatic machinery, the excess photons would lead to damage of the photosystem (Falkowski and Raven 2007). To prevent photo-oxidative damage, the cells therefore must dissipate energy before it can enter the system. This is empirically demonstrated by the increase in NPQ_NSV_ in the top ice layer, which reached values of 36.2 as F_m_’ was suppressed to near baseline values. In this high-stress zone, the “back-pressure” of electrons would have been high. Consequently, the F_v_/F_m_, even in the dark acclimated state, experienced near-collapse by day 16. Sustained F_v_/F_m_ values below 0.1 generally denotes severe, irreversible damage to the PSII reaction centers, indicating that these cells likely reached a state of terminal photoinhibiton or death (Obara et al. 2022). While the elevated NPQ_NSV_ values reflect an intense upregulation of thermal dissipation pathways designed to protect from this excess energy, the lack of a reproducible JVPII and 1/τ signal after day 16 indicates that forward electron transport was severely hindered. The persistence of measurable 1/τ kinetics up until day 16 suggests that the downstream photosystem, though heavily compromised, maintained a basal level of functionality for up to two weeks. The high NPQ_NSV_ activity thus successfully kept the photosynthetic apparatus functional for some time (likely enough for the Antarctic winter to reduce overall irradiance to non-stressful levels), thus cells may eventually survive this stress within the ice-column, even the surface, long-term.

Within the middle ice-layer, the maximum rate of electron transport (ETR_cell_) was significantly depressed compared to the bottom layer. Because electron transport downstream of the PSII is a multi-enzymatic process limited by the turnover of temperature-sensitive catalysts like RuBisCO (Young et al. 2016), the reduction of ETR_cell_ under colder temperatures indicates that the carbon fixation was constrained by a Q10 temperature effect, regardless of the photon availability. The system was also likely inhibited by a metabolic effect of hypersalinity (Kirst 1990). Osmotic stress forces the cell to redirect energy (ATP and NADPH) away from biomass production toward active ion exchange pumps and the synthesis of costly compatible solutes like proline or glycerol. This redirection of energy explains the decline in ⍰, the initial slope of the P vs. E curve, in the more saline upper and middle layers. Cells dynamically downregulate the light-harvesting efficiency as they cannot afford the metabolic costs of downstream carbon fixation under osmotic stress. This interpretation is directly validated by our ice-core melt comparison experiment on day 14. When the severe salinity stress was alleviated during processing via a direct freshwater (“FW”) melt, we observed an immediate enhancement in both E_k_ and maximum ETR_cell_ alongside a depression in ⍰ and 1/τ. This rapid kinetic divergence demonstrates that the core photosynthetic machinery was not structurally affected by the ice environment but was instead dynamically suppressed by the energetic demands of osmoregulation. Once the osmotic pressure was removed, the metabolic energy deficiency was elevated, initiating an immediate re-mobilization of photosynthetic energy toward kinetic pathways. Physiology in the under-ice water was more stable and demonstrated more efficient electron flow, a possible artifact of relatively warmer temperature, likely lower salinity than that experienced in brine channels, and lowest irradiance after attenuation through the ice.

These metabolic bottlenecks, particularly pronounced in the in the top and middle layers, reflect an “energetic debt” accumulated during long-term entrapment, which likely determines cell survival and the duration of the lag phase upon release. Importantly, the vertical physicochemical gradients across the ice column do not modulate instantaneous photosynthetic rates; they fundamentally determine how cellular energy is partitioned between growth and repair during the ice-to-water transition. Although we did not directly quantify this energetic debt within the SIC, we characterized cellular processes that point to its prolonged physiological consequences in *F. cylindrus*. In the upper ice layers, the combined stresses of high irradiance, freezing temperatures, and osmotic extremes induce chronic photoinhibition, leading to the accumulation of structural and macromolecular damage. Upon melting, these cells likely require extended recovery periods, involving large-scale transcriptional reprogramming, de novo protein synthesis, and the re-establishment of transmembrane ion gradients. Conversely, cells within the more stable, hospitable microenvironment of the bottom ice layer maintain fully functional metabolic machinery and carry reduced osmoregulatory costs, allowing them to bypass this repair phase and immediately channel post-melt captured photons into cellular division. To empirically test the link between in-ice physiological suppression and subsequent bloom potential, we used the MIC platform to quantify post-melt growth rates across distinct ice layers. However, it is important to note that these experiments were conducted in separate laboratories using different culture isolates, as logistical constraints prevented the use of identical strains across sites.

In the MICs, we observed clear differences in both incorporation into the ice matrix and post-melt viability between species, consistent with their known ecological niches. Although we lack direct photophysiological measurements to quantify in-ice stress, post-melt growth responses provide strong indirect evidence that cells experienced comparable stress regimes during entrapment. *N*. *frigida*, a known ice-associated diatom (Michel et al. 2002), showed strong incorporation into the ice column relative to the underlying water. Following melt, populations originating from the bottom ice layer exhibited higher growth rates than those from the top layer; however, cells from both layers remained viable and capable of growth. In contrast, cultures inoculated from under-ice water did not successfully establish post-melt, likely reflecting low initial cell concentrations, although reduced fitness in the water column cannot be excluded. In contrast, *P*. *glacialis*, displayed low affinity for the ice environment and a strong preference for the underlying water, consistent with its typical association with marginal ice zone and open-water conditions (Armand 1997; Scott et al. 1994; Watanabe 1988). Despite this, cultures of *P*. *glacialis* obtained from the ice were able to grow post ice-melt, with the highest growth rate observed from the bottom ice layer and no significant difference in growth rate between the top ice layer and underlying water. Cultures inoculated from the mixed culture MIC behaved similarly; these species are likely able to coexist within the ice environment due to their specific adaptations to the ice environment (*N*. *frigida*) and ice edge (*P*. *glacialis*) and do not heavily compete for resources in a way that influences post-melt growth. Considering the difference in growth rates, it is likely that out of the two species, *P*. *glacialis* contributes more significantly to initial post-melt phytoplankton blooms. Together, these results highlight how species-specific ecological strategies shape both physical incorporation into sea ice and the capacity for post-melt recovery, reinforcing the importance of vertical ice structure as a selective filter for bloom initiation potential.

The observed differences in post-melt growth between the top and bottom layers of MIC ice for both *N. frigida* and *P. glacialis* support the idea that the covariation in temperature and salinity within the ice has significant implications for ice-associated species’ bloom potential once released from the ice column. These vertical gradients do not act independently; rather, their combined effects shape the degree of physiological acclimation achieved by cells during entrapment, ultimately influencing their capacity for rapid recovery and growth upon melt.

Although post-melt growth was not directly assessed for *F. cylindrus*, the pronounced vertical structure in its physiological profiles indicates that similar constraints likely apply. The strong covariation of temperature and salinity across the ice column corresponds with clear differences in photophysiological performance, suggesting that cells in upper ice layers experience greater metabolic suppression and stress than those in the more stable bottom layers. Based on these patterns, it is likely that *F. cylindrus* populations released from the upper ice would exhibit slower recovery and longer lag phases compared to cells originating from the bottom ice, with direct implications for their contribution to early-season bloom development.

## 5. Conclusion

We demonstrate the utility of small-scale icecosm systems as a tractable and reproducible approach to investigate the incorporation, physiology and post-melt survival of several polar sea ice algal species. By employing an affordable ice tank to probe the ecophysiological responses of diverse diatom taxa to the ice environment, this study represents an initial step toward mechanistically linking physical ice properties with biological function. However, this work is only the beginning. Building on this foundation, both conceptually and experimentally, and inspired by the pioneering insights of John Raven, future icecosm studies will expand to address broader ecological, evolutionary, and ecophysiological dynamics across diverse microalgal and microbial communities. We anticipate that such efforts will deepen our understanding of how sea ice structures biological processes, ultimately advancing our ability to predict ecosystem responses in rapidly changing polar environments.

