## Supplementary Figure S1 for "An ice-bucket challenge: investigating ice algae physiology in laboratory microcosms"

1. Light quality & quantity in the SIC

The LED system used to provide irradiance to the Square Icecosm (SIC) consisted of an array of multi-wavelength LEDs. This system was controlled using an Arduino Mega to encode for a specific spectrum set to mimic the measured PAR spectrum at 5 m in clear water (Supplementary Figure S1A). The light spectrum was measured using an miniature spectrometer (Ocean Optics, FLAME-S-VIS-NIR-ES). Provided irradiance was on a 12:12-hr light:dark cycle encoded as a sinus curve to mimic changes in light intensity throughout the day (Supplementary Figure S1B). Irradiance was measured using a Universal Light Meter (WALZ, ULM-500).

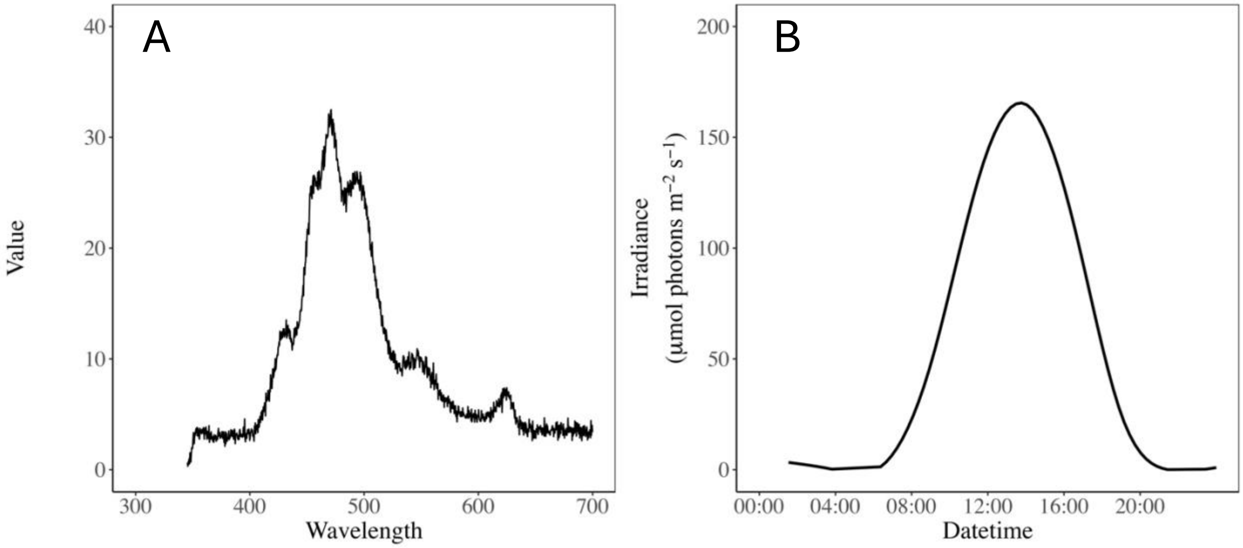


Supplementary Figure S1: (A) Measured wavelength spectrum provided in the SIC. (B) Measured daily irradiance (μmol photons $m^{-2}$ $s^{-1}$) curve in the SIC.
