## Supplementary Table S1 for "An ice-bucket challenge: investigating ice algae physiology in laboratory microcosms"

2. Experimental sampling pipeline for the SIC

Supplementary Table S1: Core types, lengths, and measurements taken each day of the SIC experiment.

| Day | Fast Repetition Rate fluorometry | Flow Cytometry |
|---|---|---|
| 0 | | x |
| 5 | | x |
| 7 | x | x |
| 9 | x | x |
| 14 | x | x |
| 16 | x | x |
| 21 | | x |
| 30 | | x |
