## Supplementary Table S2 for "An ice-bucket challenge: investigating ice algae physiology in laboratory microcosms"

3. MIC mortality analysis

Supplementary Table S2: Percentages of dead cells present in liquid cultures maintained in the light and the dark over the 4 days of Mini-icecosm (MIC) freezing and under ice water. DARK inoculum refers to a subculture of the original culture used to inoculate the MIC that was placed into darkness to compare mortality in the dark vs mortality under ice in the MIC. LIGHT inoculum refers to the original liquid culture maintained in 12:12hr light:dark conditions during freezing. Under-ice water refers to samples taken from the water underneath the MIC ice after freezing.

| Inoculum | Species | MIC | % Dead Cells |
|---|---|---|---|
| LIGHT | *N. frigida* | Mixed | 1.08 |
| | *P. glacialis* | Mixed | 7.69 |
| | *N. frigida* | Monoculture | 1.95 |
| | *P. glacialis* | Monoculture | 8.15 |
| DARK | *N. frigida* | Mixed | 3.47 |
| | *P. glacialis* | Mixed | 3.13 |
| | *N. frigida* | Monoculture | 2.26 |
| | *P. glacialis* | Monoculture | 9.48 |
| Under-ice water | *N. frigida* | Mixed | 77.85 |
| | *P. glacialis* | Mixed | 62.5 |
| | *N. frigida* | Monoculture | 71.92 |
| | *P. glacialis* | Monoculture | 35.98 |
